# Construction of distinct *k*-mer color sets via set fingerprinting

**DOI:** 10.64898/2026.02.16.706153

**Authors:** Jarno N. Alanko, Simon J. Puglisi

## Abstract

The colored de Bruijn graph model is the currently dominant paradigm for indexing large microbial reference genome datasets. In this model, each reference genome is assigned a unique *color*, typically an integer id, and each *k*-mer is associated with a *color set*, which is the set of colors of the reference genomes that contain that *k*-mer. This data structure efficiently supports a variety of *pseudoalignment* algorithms, which aim to determine the set of genomes most compatible with a query sequence.

In most applications, many distinct *k*-mers are associated with the same color set. In current indexing algorithms, color sets are typically deduplicated and compressed only at the end of index construction. As a result, the peak memory usage can greatly exceed the size of the final data structure, making index construction a bottleneck in analysis pipelines.

In this work, we present a Monte Carlo algorithm that constructs the set of distinct color sets for the *k*-mers directly in any individually compressed form. The method performs on-the-fly deduplication via incremental fingerprinting. We provide a strong bound on the error probability of the algorithm, even if the input is chosen adversarially, assuming that a source of random bits is available at run time. We show that given an SBWT index of 65,536 *S. enterica* genomes, we can enumerate and compress the distinct color sets of the genomes to 40 GiB on disk in 7 hours and 17 minutes, using only 14 GiB of RAM and no temporary disk space, with an error probability of at most 2^−82^.

## 1 Introduction

Indexing large collections of microbial reference genomes to support similarity search is now a widespread task in modern genomics, with the the colored de Bruijn graph model currently dominant paradigm [4, 12, 14]. In this model, each reference genome is assigned a unique color, and each *k*-mer is associated with a *color set*, that is, the set of colors of the references that contain that *k*-mer.

Early incarnations of this model used a binary matrix to represent color information [16, 19], where rows correspond to the k-mers of the dataset and the columns to colors (genomes). Cell [*i, j*] of the matrix is 1 if and only if the *k*-mer (with rank) *i* occurs in genome *j*. Rows of the matrix correspond to color sets. When stored naively, the matrix representation is large—a collection of 64 thousand Salmonella genomes results in a matrix of some 44 trillion cells. However, a particular property of the color matrix, especially manifest by pangenomes, is that many rows (i.e., color sets) are identical, a phenomenon borne out of the co-occurrence of *k*-mers in genomes. Identification and removal of duplicate rows is thus a key step in reducing space. Another important space-saving technique is to judiciously choose the representation used for individual color sets (those that remain after duplicate detection) according to the density of the color set, an approach borrowed from the search engine literature [22]. Another important reason to compress the matrix row-wise (as opposed to column-wise) is that important applications, such as pseudoalignment, which involve intersection or thresholded-union of colors sets, require fast access to *rows* of the matrix.

Ideally, we would like to arrive at a row-compressed representation of the matrix without having to realise the whole uncompressed matrix as an intermediate step. Various authors have made attempts at efficient (compressed) matrix construction. Metagraph [17] constructs the matrix column-wise, which is easy to implement, but loses the ability to deduplicate (or otherwise compress) rows until the whole matrix is in hand, thus requiring a huge amount of intermediate space. Another approach (adopted by, e.g., Bifrost [14], GGCAT [8]) is to use dynamic set data structures to construct the rows incrementally, but this makes parallelism complicated, and still makes the intermediate space significantly higher than the final space. Current methods deduplicate rows of *k*-mers that are in the same unitig, but deduplicating across unitigs during construction remains challenging.

In this paper, we present an algorithm that (1) parallelizes well, requiring only a certain well-supported set of atomic CPU instructions, and no higher-level synchronization primitives (2) deduplicates rows during construction, even across unitigs (3) does not require dynamic data structures: instead, a static data structure is constructed in multiple stages. We describe a Monte Carlo algorithm that constructs the set of distinct color sets for the *k*-mers directly in any individually compressed form, using the unitigs of each reference genome. The method performs on-the-fly deduplication via incremental fingerprinting. We provide a strong bound on the error probability of the algorithm, even if the input is chosen adversarially, assuming that a source of random bits is available at run time.

We show that given an SBWT index [3] of 65,536 *S. enterica* genomes, we can enumerate and compress the distinct color sets of the genomes to 40 GiB on disk in 7 hours and 17 minutes, using only 14 GiB of RAM and no temporary disk space. For context, the original genome files in FASTA format occupy 294 GiB on disk.

The remainder of this paper is structured as follows. In the next section we lay down notation and basic concepts used throughout. Then, in Section 3, we describe the main tenents of our approach, including details for its efficient implementation. In Section 4 we describe results of extensive experiments measuring performance. Conclusions and avenues for future work are then offered.

## 2 Notation and preliminaries

### Strings, *k*-mers and color sets

A string of length *n* is an array *T*[0..*n* − 1] with elements from an alphabet Σ. We use both closed and half-open interval notations for substrings, that is, the substring of *T* starting from index *i* and ending at index *j*, not including index *j*, is denoted with *T*[*i*..*j*) or *T*[*i*..*j* − 1]. In this paper, we assume the DNA alphabet Σ = {*A, C, G, T*}, though our algorithm generalizes to any alphabet. A *k-mer* is a string of length *k*. In the context of DNA, a *k*-mer is called *canonical* if it is lexicographically less or equal to its reverse complement on the opposite strand. The *k-spectrum* of a string *T*, denoted with *S*_*k*_(*T*), is the set of distinct *k*-mers contained in *T*, that is, *S*_*k*_(*T*) = {*T*[*i*..*i* + *k*) | 0 ≤ *i* < |*T* | − *k* + 1}. The input to *S*_*k*_ is also allowed to be a *list* of strings. For a list of strings *T*_0_, …, *T*_*m*−1_, we define 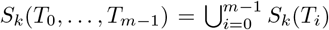. Given a list of strings *T*_0_, …, *T*_*m*−1_, the *color set* of a *k*-mer *x* ∈ *S*_*k*_(*T*_0_, …, *T*_*m*−1_), denoted with cs(*x, T*_0_, …, *T*_*m*−1_), is the set of indices of input strings that contain *x*, that is, cs(*x, T*_0_, …, *T*_*m*−1_) = {*i* ∈ {0, …, *m* − 1} | *x* is a substring of *T*_*i*_}. When the input strings *T*_0_, …, *T*_*m*−1_ are clear from the context, we drop them in the notation and write just cs(*x*).

### De Bruijn Graphs

The de Bruijn graph of a set of *k*-mers is a graph where each distinct *k*-mer in the set is a node, and there is an edge from *k*-mer *x* to *k*-mer *y* iff *x*[1..*k*) = *y*[0..*k* − 1). The de Bruijn graph of a list of strings *T*_0_, …, *T*_*m*−1_ is the de Bruijn graph of their *k*-mers *S*_*k*_(*T*_0_, …, *T*_*m*−1_). A *unitig* is a maximal non-branching simple path (*v*_0_, …, *v*_*ℓ*−1_) in the de Bruijn graph in the following sense: for *i* > 0, the in-degree of *v*_*i*_ is 1, and for *i* < *ℓ*− 1, the out-degree of *v*_*i*_ is 1, and the path cannot be extended in either direction without breaking these properties.

### Rank queries

Given an array of bits *B*[0..*n*), a rank-1 query at position *i*, denoted with *rank*_1_(*B, i*), counts the number of 1-bits in *B*[0..*i*). Likewise, a rank-0 query, denoted with *rank*_0_(*B, i*) counts the number of zero-bits in *B*[0..*i*). When we refer to just “rank queries,” or omit the subscript 1 from the notation, we mean rank-1 queries. There exist practical data structures to support both types of rank queries on *B* in *O*(1) time using *o*(*n*) bits of extra space [20].

### 2.1 State of the art in compressed color set construction

Most currently available tools build an uncompressed color set representation before compressing the data into the final form. For example, the recommended construction pipeline of Metagraph [17] is to first index the distinct *k*-mers without colors, then construct the color annotation of *k*-mers of each color independently, then combine the annotations and finally compress them using a variety of methods [18, 9, 5]. While the final space can be very small, the intermediate space during construction in memory or on disk can be orders of magnitude larger.

**Bifrost** [14] first constructs the unitigs of the input, indexes them with minimizer hashing, and then employs dynamic set data structures to construct the color sets incrementally. Space is saved by constructing the coloring at the level of *subunitigs*, that is, instead of associating *k*-mers to color sets, Bifrost breaks unitigs into runs of *k*-mers whose color set is the same, and colors the runs instead of individual *k*-mers. This provides color set deduplication within unitigs, but not *across* unitigs. The encoding of each color set is chosen to exploit properties like density and long runs of consecutive color identifiers. Parallel synchronization is managed with a system of mutex locks: instead of a single global lock, the data structure is striped with many locks so that each lock only locks one slice.

The **GGCAT** tool [8], which is at the heart of the color construction pipelines of Themisto [4] and Fulgor [12], constructs both the unitigs and the color sets of the *k*-mers at the same time by bucketing *k*-mers by minimizers and assembling to unitigs, and tracking color information at the same time. The color sets in unitig fragments are maintained as dynamic resizable vectors of integers encoded with a custom variable-length encoding inspired by Google Varint. This also provides color set deduplication within unitig fragments, but not across unitigs until the end. Finished color sets are inserted into a global hash table, and compressed using Lempel-Ziv encoding combined with Huffman coding. The intermediate working space remains significantly larger than the final compressed space.

In general, using dynamic set data structures for this problem is not ideal since their space usage is usually significantly higher than their static counterparts, and they lead to memory fragmentation due to frequent reallocations. GGCAT 2.0 uses a custom dynamic memory allocator to mitigate this issue to some extent, but it can still use up to twice as much memory as needed. Another drawback of dynamic data structures is that they do not work well with parallelism. A synchronization mechanism is usually needed to prevent two threads from modifying the same structure at the same time, which results in a communication overhead between threads and wasted time due to threads having to wait for their turn to access the data.

### 2.2 Sparse-dense color set representation

The goal of our work is to build a compact representation of the distinct color sets of the *k*-mers of the input genomes. There is a growing literature on ways to compress the color sets (see [6] and references therein), much of it stemming from literature on inverted index compression [22]. In this work, our target is the representation from Themisto [4] (Fulgor [12] and Bifrost [14] employ similar schemes). Since our algorithm depends on details of the representation, we now recall the details of the representation here.

Let *d* be the number of distinct color sets. Each color set is encoded either in sparse or dense form, depending on which representation requires less space (this is defined precisely in a moment). Let *B* be a bit vector of length *d* such that *B*[*i*] = 1 if color set *i* is stored in sparse form, and *B*[*i*] = 0 if it is stored in dense form. The bit vector *B* is indexed for rank queries. A sparse set is stored as a sorted list of its elements, encoded using ⌈log *m*⌉ bits per element. The representations of all sparse sets are concatenated into a single array *A* of ⌈log *m*⌉ -bit integers. The starting points of each set are stored in an offset array *O* of 64-bit integers, where *O*[*j*] gives the starting position of the *j*-th sparse set. A sparse set of *s* elements therefore occupies ⌈log *m*⌉*s* + 64 bits in total. Dense color sets are stored as bitmaps of length *m* bits. All the bitmaps are concatenated into a single bitmap *D*. Since all dense sets have the same size, no offset array is required, and each dense set occupies exactly *m* bits. A color set of size *s* is stored in sparse form if ⌈log *m*⌉*s* + 64 *< m*, and in dense form otherwise. To access color set *i*, we first inspect *B*[*i*]. If *B*[*i*] = 1, then its rank among sparse sets is *j* = *rank*_1_(*B, i*), and its elements are stored starting at offset *O*[*j*] in *A*. If *B*[*i*] = 0, then its rank among dense sets is *r* = *rank*_0_(*B, i*), and its bitmap starts at bit index *rm* in *D*.

## 3 Algorithm

### 3.1 Problem statement and setup

To simplify our exposition, we model a genome as a *single* string from the DNA alphabet. The input to our algorithm is a set of *m* genomes 𝒢 = {*G*_0_, …, *G*_*m*−1_}. Multi-string genomes can be easily supported in implementation. The goal is to construct the set of distinct color sets of 𝒢, and an index structure that can quickly return the color set of any query *k*-mer.

We assume we have data structure support for de Bruijn graph operations to (1) compute the outdegree of a node (2) compute the indegree of a node (3) move from a node to any of its in-neighbors. These can be provided with any *k*-mer lookup structure, e.g. with the SBWT or the Sshash index, or any hashing-based compact de Bruijn graph data structure.

Further, we assume that we have an injective (i.e. perfect) hash function *h* : *S*_*k*_(𝒢) → {0, …, *n* − 1} mapping the distinct *k*-mers to distinct integers. The largest hash value *n* − 1 is allowed to be greater than |*S*_*k*_(𝒢)|, but should be reasonably small to obtain good space efficiency. Examples of such functions include Sshash [21], which can be used to map a *k*-mer to its starting position in the unitigs of the input genomes, and SBWT [3], which can be used to map each *k*-mer to its *colexicographic rank* among the input *k*-mers (roughly speaking). Any generic perfect hash function, e.g. PtrHash [13] or Recsplit [11], will also do.

### 3.2 Algorithm

We introduce the notion of a *color-set covering k-mer set*. Given a list of genomes 𝒢 = *G*_0_, …, *G*_*m*−1_ (strings), a color set covering *k*-mer set is a set of *k*-mers *A* ⊆ *S*_*k*_(𝒢) such that every color set is found at least once. More formally:

#### Definition 1.

***(Color-set covering*** *k****-mer set)***

*Given a set of strings* 𝒢 = *G*_0_, …, *G*_*m*−1_, *a set of k-mers A* ⊆ *S*_*k*_(𝒢) *is a color-set covering k-mer set if for every x* ∈ *S*_*k*_(𝒢) *there exists a k-mer y* ∈ *A with* cs(*x,* 𝒢) = cs(*y,* 𝒢).

Finding a small color-set covering *k*-mer set quickly is a major piece of our approach. We proceed in three phases. **In the first phase**, we find a color-set covering *k*-mer set *C*_1_ ⊆ *S*_*k*_(𝒢), which may not be minimal, but tends to be much smaller than *S*_*k*_(𝒢) because it exploits the property that in genomic data, the color sets of *k*-mers in the same unitig tend to be the same. We call the *k*-mers in *C*_1_ *key k-mers*. **In the second phase**, we use a fingerprinting scheme to shrink this set to obtain a *minimal* color-set covering *k*-mer set *C*_2_ ⊆ *C*_1_, containing exactly one *k*-mer per distinct color set in the data, called the *sufficient k-mers*. **In the third phase**, we construct the color sets of *k*-mers in *C*_2_ into a sparse-dense structure as defined in Section 2.2 See Figure 1 for a visual representation of the inputs and outputs to each phase. We now proceed to describe the phases in detail.

**Figure 1:**
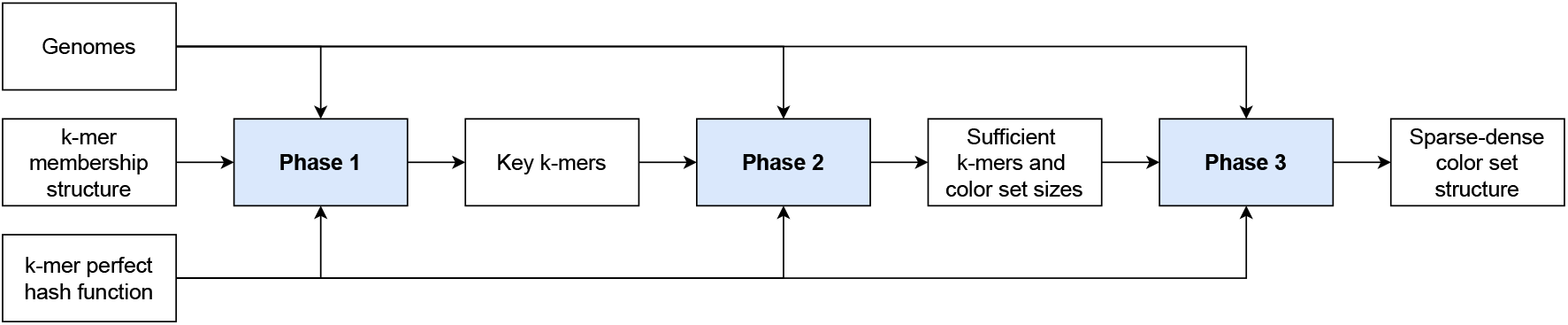
Algorithm overview. The *k*-mer membership structure can be discarded after Phase 1.

**Figure 2:**
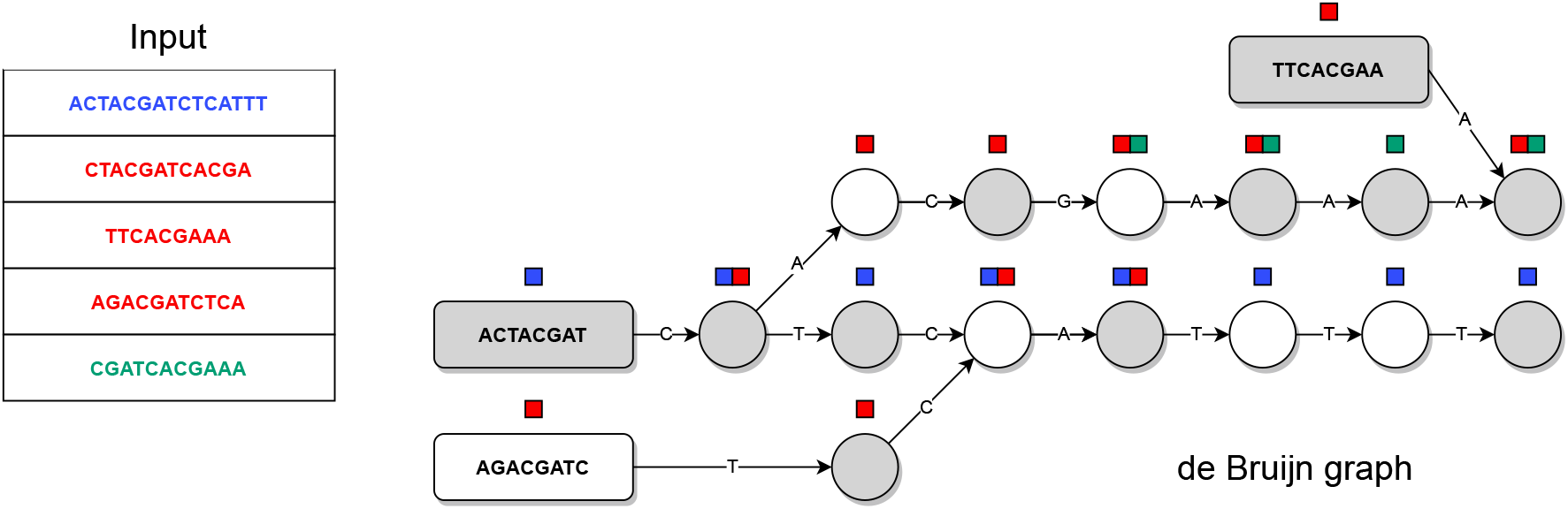
Phase 1. Each node represents a *k*-mer, with *k* = 8. To keep the figure clean, the *k*-mer labels are written only for source nodes. The edge labels are the last characters of the destination *k*-mers. So-called *key k*-mers are the nodes with a gray background. The color sets of the nodes are shown above the nodes. The color set of each white node can be obtained by walking forward to the nearest gray node.

#### Phase 1: Marking key *k*-mers

In this phase, we identify a set of *key k-mers*, such that every distinct color set is the color set of at least one key *k*-mer. A *k*-mer *x* is key if any of the below holds:

1. *x* is the last *k*-mer of a string in the input genomes.
2. *x* has an out-neighbor that is the first *k*-mer of a string in the input genomes.
3. *x* is the last *k*-mer of a unitig of the de Bruijn graph of all the *k*-mers, that is, *x* has outdegree not equal to 1, or *x* has an out-neighbor with in-degree greater than 1.

We initialize a bit vector of length *n* to mark the hash values of *k*-mers according to the hash function *h* that match any of the three above conditions. To mark *k*-mers matching conditions 1 or 2, we read through the input genomes and apply *h* on the last *k*-mer of each sequence, and the de Bruijn graph in-neighbors of the first k-mer of each sequence. Then, we use the de Bruijn graph structure to identify and mark all *k*-mers fulfilling condition 3.

If *x* is not marked, it has exactly one out-neighbor *y*, and, moreover, *y* has the same color set as *x* (See Lemma 1 in [4]). We call *y* the successor of *x* and denote *y* = `succ`(*x*). Let `depth`(*x*) be the smallest *ℓ* such that `succ`^*ℓ*^(*x*) is marked, where `succ`^*ℓ*^(*x*) is the successor function iterated *ℓ* times. Our construction guarantees that such *ℓ* always exists, as shown by the following lemma.

##### Lemma 1.

*Every k-mer x has a finite* `depth`(*x*).

*Proof*. For any marked *k*-mer *x*, we have `depth`(x) = 0. If a node is not marked, it has a unique successor. We need to show that iteratively applying `succ`(*x*) eventually leads to a marked node. For a contradiction, let us assume this does not happen. Then, the sequence of successors eventually gets stuck in a cycle of non-marked nodes. By condition 3, all nodes on the cycle have both in-degree and out-degree 1. This means that there must exist a marked node on the cycle because the *k*-mers on the cycle come from some original sequence, whose last *k*-mer is marked, and that *k*-mer is somewhere on the cycle.

##### Lemma 2.

*For every distinct color set in the input, there is at least one marked k-mer that has that color set*.

*Proof*. Induction on `depth`(*x*). Suppose the color sets of all *k*-mers with `depth`(*x*) ≤ *ℓ* are covered, and let *f*(*x*) be any key *k*-mer whose color set is the same as that of *x*. For the base case, at *ℓ* = 0 the claim holds because nodes with depth 0 are marked, and we can set *f*(*x*) = *x*. Consider *k*-mers with `depth`(*x*) = *ℓ* + 1. Since `depth`(*x*) > 0, they each have a unique successor, whose color set is cs(*f* (`succ`(*x*))), which is correct by induction since `depth`(`succ`(*x*)) = *ℓ*.

#### Phase 2: Computing fingerprints and marking sufficient *k*-mers

In this phase, we compute a fingerprint of the color set of every key *k*-mer marked in the first phase, and in the end, assign ids 0, 1 … to distinct fingerprints. The input is the key *k*-mer mark bit vector from Phase 1, the input genomes 𝒢, and a perfect hash function on the *k*-mers of 𝒢. The output is a bitvector marking sufficient *k*-mers, and an array containing the sizes of the color sets of the sufficient *k*-mers in the order they appear in the bitvectors.

The fingerprinting scheme is an instance of *tabulation hashing*, a generic technique dating back to 1970, originally used for fingerprinting distinct chess positions [23]. First, we assign a random *ℓ*-bit fingerprint for each distinct individual color, and store these in a table. The higher the value of *ℓ*, the smaller the probability of error in the algorithm will be. We denote the chosen fingerprint of color *c* with *f*(*c*). The fingerprint of a *set* of colors will be the xor of the fingerprints of the individual colors in the set. We denote the fingerprint of set *A* with *F*(*A*), that is, *F*(*A*) = ⊕_*c* ∈ *A*_ *f*(*c*). This manner of fingerprinting lends itself to very clean collision analysis. The next lemma shows that *F* is a universal hash function family over the random choice of *f*.

##### Lemma 3

*Let A and B be two distinct sets of colors and select the function f uniformly at random from all functions* 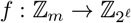, *where m is the total number of possible colors. We have* Pr(*F*(*A*) = *F*(*B*)) = 2^−*ℓ*^.

*Proof*. Since *A* ≠ *B*, there exists a color *c* that occurs in exactly one of *A* and *B*. Without loss of generality, suppose that *c* ∈ *A* and *c* ∉ *B*. We have that *F*(*A*) = *F*(*B*) if and only if *F*(*A*\ {*c*}) ⊕*f*(*c*) = *F*(*B*), where ⊕ is the xor operation. Rearranging, this means *f*(*c*) = *F*(*A*\ {*c*}) ⊕*F*(*B*). There are 2^*ℓm*^ possible functions *f*. To count the functions for which the equation holds, first choose a value for *f*(*x*) for each *x* ≠ *c*. There are 2^*ℓ*(*m* −1)^ possibilities. This determines the value of *F*(*A* \ {*c*}) ⊕*F*(*B*), so there is exactly one way to choose *f*(*c*) to make the equation hold. Hence, the fraction of functions for which the equation holds is 2^*ℓ*(*m* −1)^/2^*ℓm*^ = 2^−*ℓ*^. □

This gives us the following bound on the probability of a hash collision across all given color sets.

##### Lemma 4.

*Given a set of distinct sets A*_0_, …, *A*_*N* −1_, *the probability that there exists two sets A*_*i*_ ≠ *A*_*j*_ *such that F*(*A*_*i*_) = *F*(*A*_*j*_) *is at most N* ^2^*/*2^*ℓ*+1^.

*Proof*. By the union bound, and Lemma 3, the probability is at most 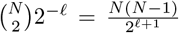. Rounding the numerator up to *N* ^2^ gives the claimed bound. □

For example, for *ℓ* = 128 and *N* = 10^9^, we have a collision probability of at most 10^18^/2^129^ ≈1.47 · 10^−21^. This manner of fingerprinting has the useful property that the fingerprint of a set can be built incrementally by initializing the fingerprint to a *ℓ*-bit string of zeroes, and individually xorring the fingerprints of the set’s colors in any order.

To build the fingerprints, we initialize an array *A*[0, *s* −1] of *ℓ*-bit integers, where *s* is the number of key *k*-mers identified in the first phase. We preprocess the key *k*-mer marks bit vector *B* from Phase 1 for rank queries, so that the fingerprint of *k*-mer *x* is at *A*[`rank`(*B, h*(*x*))]. Then, we iterate over all the *k*-mers in each input genome, and xor the fingerprints of the colors of each genome to the corresponding indexes in *A*. See Figure 3.

**Figure 3:**
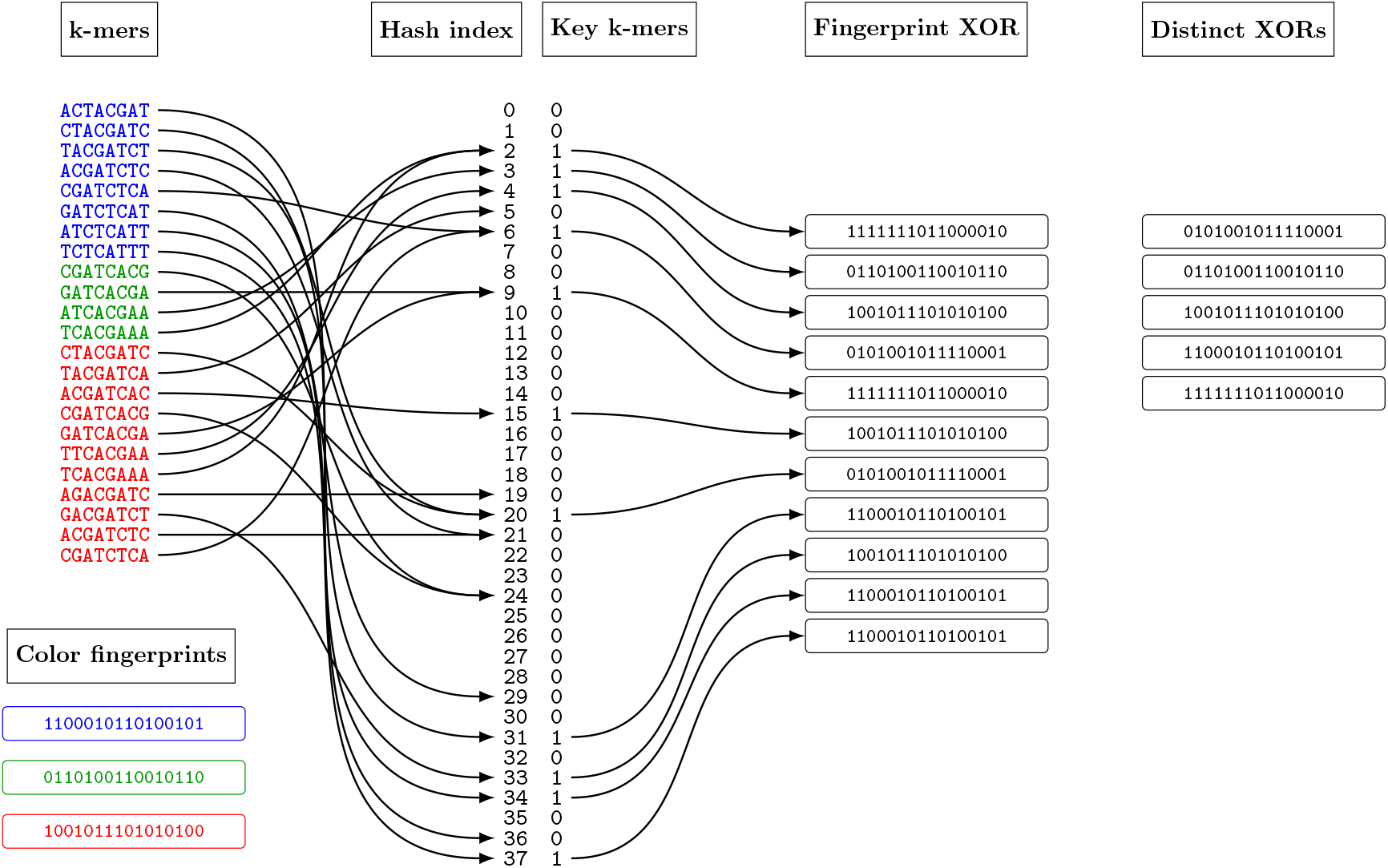
Fingerprinting in Phase 2. At the start of the phase, each genome (color) is associated with a random *ℓ*-bit fingerprint. In this picture, *ℓ* = 16. Each *k*-mer is hashed with the perfect hash function, and in the event it turns out to be a key *k*-mer, the fingerprint of the color is xorred into the aggregate fingerprint of the set. In the end, the aggregate fingerprints of the sets are sorted and deduplicated.

This can be done in parallel using atomic instructions to apply the xor-operations to elements of *A*. Since xor is commutative, no further synchronization between threads is required because any order of updates results in the same final result.

At the same time as building *A*, we also build another array *E* of the same length, where *E*[*i*] stores the number of distinct elements xorred into *A*. The arrays *E* can also be built in parallel with widely supported atomic increment operations.

We obtain the set of distinct fingerprints by sorting and deduplicating in-place. Let *d* be the number of distinct fingerprints found. The set of distinct fingerprints are assigned ids from 0, …, *d* − 1. We pick for each distinct color set a single representative *k*-mer that has that color set. The representative among *k*-mers that have a given color set is defined as the one which has the smallest hash value according to *h*. The representatives form the set of sufficient *k*-mers for this dataset. We mark these *k*-mers in a bit vector *B*_2_ of length *n*, and index the bit vector for rank queries.

In the final phase, we build the color sets of the representative *k*-mers.

#### Phase 3: Building the sparse-dense structure

In this phase, we build the sparse-dense structure described in Section 2.2 for the sufficient *k*-mers marked in the previous phase. Let *B*_2_ be the bit vector marking the sufficient *k*-mers from the previous phase. We start by allocating a “skeleton” of the final sparse-dense data structure, containing the bit vector that marks sparse sets, the space for the concatenation of dense bitmaps and sparse integer lists, and the offset array, all described in Section 2.2. The marking bit vector is allocated and populated first based on the sizes of the sets, which informs us how many bits are required for the sparse and dense components. To fill in the elements of the sets, we run one final pass over the input genomes, applying hash function *h* on each *k*-mer *x*, and whenever *B*_2_[*h*(*x*)] = 1, we query the skeleton to find where the set is stored in the data structure, and add the element to the set. Some care is required to append to the same sets from multiple threads. These details are explained toward the end of the next subsection.

### 3.3 Implementation details and space analysis

#### Phase 1

Let *n* be the number of distinct *k*-mers in the input. Phase 1 requires a set membership structure on the *k*-mers in order to measure indegrees and outdegrees of *k*-mers in the de Bruijn graph, and to traverse to neighboring *k*-mers. We also require a perfect hash function on the *k*-mers. The SBWT [3] and the Sshash [21] data structures both double as a *k*-mer membership structure and a perfect hash function: for SBWT, the hash value is the position of the *k*-mer on the SBWT, and in Sshash it is the starting point of the *k*-mer in the concatenation of the unitigs of the data. These data structures typically take 2.5 −15 bits per *k*-mer on real-world sequencing data [3, 21]. In the worst case, up to *k* log *σ* bits are required, where *σ* is the size of the alphabet—this represents the case where the *k*-mers do not overlap at all, so we cannot do better than encoding them individually using log *σ* bits per character. The key *k*-mer marks take another *n* bits, where *n* is the size of the range of the perfect hash function. In summary, the space used in phase 1 is kmers + *n*, where kmers is the combined space of the *k*-mer membership structure and the perfect hash function into {0, …, *n* − 1}, with practical values ranging from 3.5 − 16 bits per distinct *k*-mer. In this work, we use the SBWT has a perfect hash function, but we emphasize this is not a fixed choice.

#### Phase 2

In Phase 2, we retain the key *k*-mer bit vector *B*, but we no longer need the set membership structure of *k*-mers, only the perfect hash function. Let *n*_*key*_ be the number of key *k*-mers and *ℓ* be the fingerprint length in bits. Since each unitig introduces only at most one key *k*-mer, and each input sequence introduces at most two, we can bound *n*_*key*_ ≤ *u* + 2*m*, where *u* is the number of unitigs and *m* is the number of input sequences. The fingerprint array takes *n*_*key*_*ℓ*bits. To deduplicate the fingerprints in the end and assign ids and sizes to unique color set ids, we build and sort an array of triples (key *k*-mer hash value, set fingerprint, set size), using the set fingerprint as the primary sort key, breaking ties using the key *k*-mer hash value. The fingerprints take *ℓ* bits each, the hash value takes ⌈log_2_ *n*⌉ bits and the set size ⌈log_2_ *m*⌉ bits, so in total, the triples take *n*_*key*_(*ℓ* + ⌈log_2_ *n*⌉ + ⌈log_2_ *m*⌉) bits, (though in practice to simplify code, we use 64-bit integers for the second and third elements of triples). We assign integer identifiers 0, 1, … for the distinct set fingerprints in the sorted order. The smallest key *k*-mer that has each distinct fingerprint is chosen as the representative *k*-mer for that color set. These are marked in the bit vector *B*_2_ of length *n*. In summary, the space needed for phase 2 is `PHF` + *n*_*key*_(*ℓ* + ⌈log_2_ *n*⌉ + ⌈log_2_ *m*⌉) + 2*n* + *o*(*n*), with *n*_*key*_ ≤ *u* + 2*m*, where `PHF` is the size of the perfect *k*-mer hash function, the *o*(*n*) term is for the rank structures on the key *k*-mer and sufficient *k*-mer marks.

#### Phase 3

We retain from the previous phase the perfect hash function, the bit vector *B*_2_ marking sufficient *k*-mers, and the array of color set sizes. We build the sparse-dense color set representation described in Section 2.2 directly. Since we know the size of each distinct set, we are able to pre-allocate all the required memory: the bit vector that marks sparse sets, the concatenation of the sparse sets, the offset array to the sparse sets, and the concatenation of bitmaps of dense sets. All this takes ⌈log *m*⌉*s* + 64*d*_*sparse*_ + *d*_*dense*_*m* + *d* + *o*(*d*) bits, where *d* is the number of distinct color sets, *d*_*sparse*_ is the number of sparse sets and *d*_*dense*_ the number of dense sets (*d*_*sparse*_ + *d*_*dense*_ = *d*), and *s* is the total number of elements in all sparse sets. The overhead on top of this is the size of the perfect hash function, the 64-bit color set sizes array and the sufficient *k*-mers bit vector, for a total overhead of `PHF` + 64*d* + *n* + *o*(*n*).

#### Parallelism in phase 3

In this phase, we build the sets by adding elements one by one, but this makes parallelism nontrivial since two threads might try to add elements to the same set at once. Incidentally, the construction algorithm in Bifrost [14] runs into the same issue, which is solved there by using mutexes to ensure that no two threads operate on the same set at once. Our construction method and color set representation allows for a completely lock-free solution. Updating the dense bitmaps in parallel is simple since modern CPUs support atomically setting a single bit in a word. Updating sparse sets is more tricky because each thread needs to *append* to the same list. However this can be handled by storing the offset where the next item should be stored, using an atomic fetch-increment operation to read the offset, and then write to the fetched offset. The atomic fetch-increment ensures that no two threads append to the same position. The write itself is also tricky since we are not operating on full memory words but rather ⌈log *m*⌉ bit integers. However the write can be achieved atomically by zeroing the target bit region with atomic and-operations, and writing the bits with atomic or-operations — no two different threads ever modify the same bits at once, so all interleavings of the parallel operations will result in the same final result.

#### Running from input genomes with repeated *k*-mers

The fingerprinting assumes that every color fingerprint is xorred into the aggregate color set fingerprint only once. Otherwise, if an element is xorred an even number of times, it is the same as not xorring at all, and the collision analysis in Lemma 3 breaks. There is no easy fix for this, since if we could detect whether a given color has been added to a given set, it would amount to having a dynamic set membership data structure for each of the color sets being constructed, which is difficult to implement in small space — in fact, this type of a scheme is precisely what we aim to avoid with the fingerprinting. Instead, when running from genomes that might have duplicate *k*-mers, we add a deduplicating layer before xorring to avoid xorring the same color twice. We implement this by maintaining the set of perfect hash values of *k*-mers that have been processed so far for the current genome. This set is stored as a hash table, until the space of the hash table becomes bigger than *n*, at which point we switch to an indicator bit vector of length *n* marking the perfect hash values of those *k*-mers that have been processed. Each deduplicating structure is maintained in memory only for the duration of the processing of its corresponding genome.

#### Constructing directly to disk

To decrease the peak working space, we can construct the final sparse-dense data structure directly on disk. This is possible because after phase 2, we know in advance the size and the preferred representation of each set. We can thus allocate all the space for the final data structure on disk up front. Then, we split the input genomes into *p* ≤ *m* chunks (e.g. *p* = 8), and process each chunk of genomes in turn. Chunk *i* will contain the colors in the range [*i*⌈*m*/*p*⌉ .. min((*i* + 1) ⌈*m*/*p*⌉, *m*)). For each chunk, we run phase 3 to build a sparse-dense structure for the colors of the chunk, and then stream over the file on disk, writing the constructed colors to their memory locations in the final data structure. This way, only a fraction of approximately 1/*p* of the final sparse-dense data structure is held in memory at once. Overall, this means running *p* passes over the output file, so *p* should be kept reasonably small.

## 4 Experiments

We implemented our method using an SBWT index to provide both *k*-mer lookup and the perfect hash function for the input *k*-mers: the hash value of a *k*-mer is its position in the SBWT. The queries are sped up by utilizing the longest common suffix array [1] to enable streaming *k*-mer lookup via the *k*-bounded streaming matching statistics algorithm [2]. Both DNA strands are indexed, and these data structures take roughly 10 bits per *k*-mer (20 bits per canonical *k*-mer).

### 4.1 Hardware

All experiments were conducted on a single Linux server running Ubuntu 20.04 with kernel 5.15.0-153-generic. The system is equipped with 504 GiB of available RAM and an AMD Ryzen Threadripper PRO 3975WX processor (32 cores, up to 64 threads, 3.5 GHz clock, 32 × 32 KiB L1 cache, 32 × 512 KiB L2 cache, 128MiB L3 cache). The storage consists of four 4 TB Seagate ST4000NM000A CMR HDDs, with observed^1^ sustained write speeds of approximately 400 MiB/s.

### 4.2 Datasets

We set *k* = 31 for all experiments.

- Salmonella: Randomly selected 2^16^ = 65536 *Salmonella enterica* genomes from the AllTheBacteria [15] dataset.
- Random: Randomly selected 2^14^ = 16384 genomes from the AllTheBacteria [15] dataset.

We argue that these datasets represent the important performance extremes in the indexing problem at hand. The Salmonella dataset represents a low-diversity use case with large color sets, and Random represents a high-diversity use case with small color sets. To study scaling of the methods, we run them on random subsets of 2, 4, 8, 16, … genomes. Various key statistics on the datasets are listed in Table 2.

### 4.3 Baselines

We compare our method against methods from the literature that are able to construct the color sets directly without first constructing an uncompressed representation. The competing methods are as follows:

- **GGCAT 2** [8] (commit id `14e8159`). The construction algorithm in GGCAT 2 is a highly optimized minimizer bucketing method, which combines unitig construction and color indexing at the same time.
- **Bifrost** [14] (commit id `b7659dd`). The algorithm first assembles the *k*-mers into a de Bruijn graph with minimizer hashing, and constructs the sets incrementally with dynamic data structures.

See Section 2.1 for a more detailed description of the inner workings of GGCAT 2 and Bifrost. We work under the assumption that the *k*-mers have already been indexed, so we do not include the time and space for *k*-mer indexing in our results, except for GGCAT 2, which merges the coloring and *k*-mer indexing phases in a way that they can not be separated from one another. In case of Bifrost, we modified the source code to add a timer that starts from the moment when the coloring process starts, and ends when the coloring is written to disk. In both full datasets Salmonella and Random, the peak memory of Bifrost was observed to occur during color set construction, so no finer-grained memory measurement was needed.

We omit the pseudoalignment tools Fulgor [12] and Themisto [4] because their color construction algorithms are almost entirely outsourced to GGCAT. We also omit Metagraph [17] because the recommended way to construct the color sets is by first enumerating *k*-mer sets of colors (instead of color sets of *k*-mers), then transposing the relationship to obtain the color sets. This workflow blows up the intermediate space, because it is unable to compress the color information across genomes, where most of the redundancy lies, until later stages in the pipeline. Metagraph also provides a command-line option to construct directly in the final row-major form, but this does not appear to effectively exploit cross-genome redundancies until the end. For example, running annotation with options`--anno-type row, --fwd-and-reverse` and`--mem-cap 1gb` consumed 53 GiB of RAM already on 2048 Salmonella genomes, whereas our tool requires only 3 GiB. There are various other command line options and alternative workflows that could be employed, but designing such workflows requires substantial expertise from the user and knowledge of the structure of the input data, and showcasing a potentially sub-optimal workflow risks making Metagraph look worse than it is capable of, and thus we leave this out of scope of the present experimental study.

### 4.4 Results

Our method was run both fully in-memory, and in the mode where we construct the final sparse-dense structure directly to disk in *8 pieces* as described in Section 3.3. Time, memory and disk space of the final output for all benchmarked tools are listed in Table 1. Figure 6 shows time and memory plotted as the function of the number of genomes, and Figure 4 shows the scaling with respect to the number of threads.

**Table 1:**
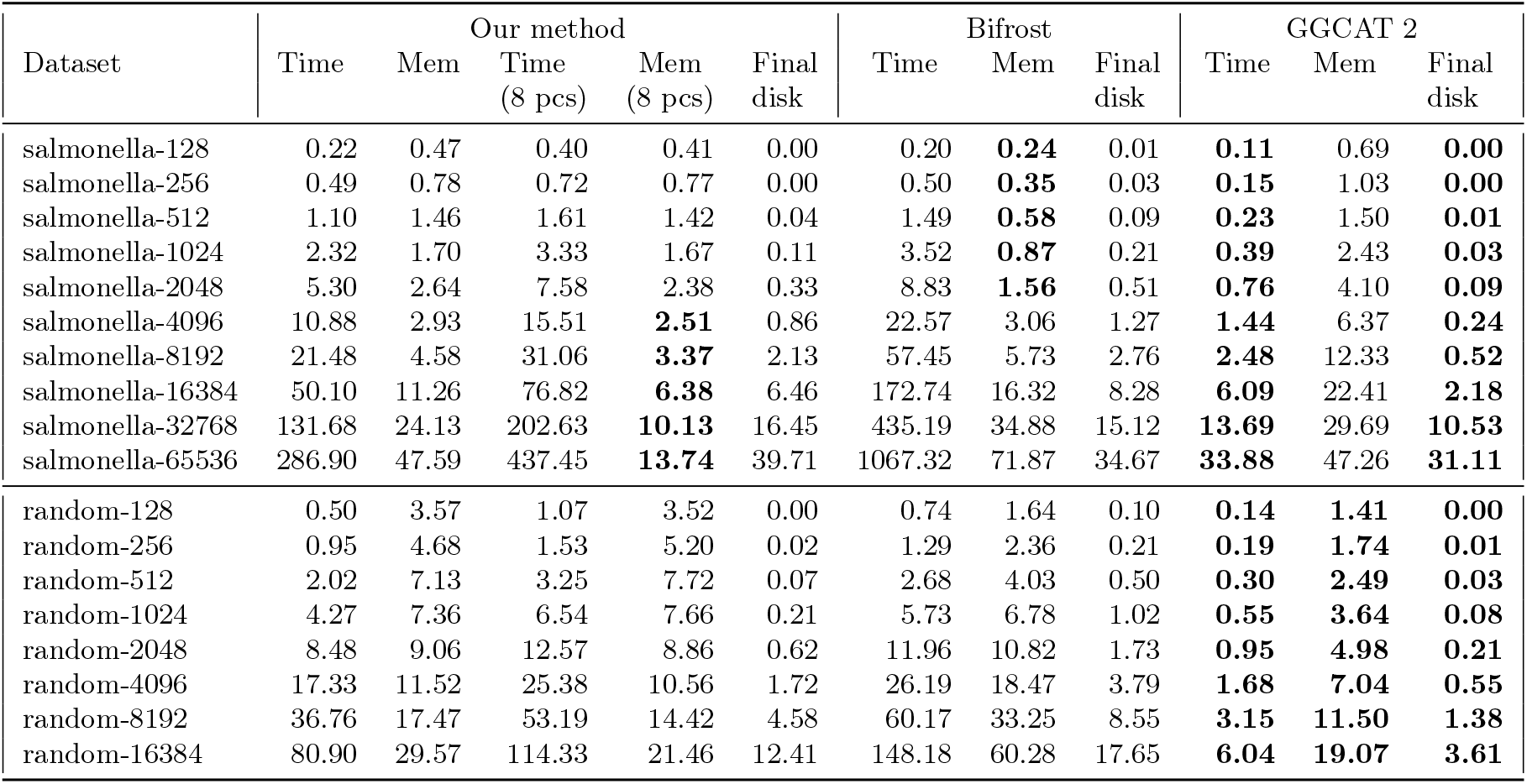
Performance metrics for *n* ≥ 128 genomes. The unit of time is minutes, the unit of memory and final size on disk is gigabytes. Time and memory for our method is reported both when running fully in-memory, and when writing the final color sets to disk in 8 pieces (8 pcs). For all tools, the final disk includes only the coloring part of the data structure. The lowest time, memory and final disk values in each row are bolded.

| Dataset | Our method |  |  |  |  | Bifrost |  |  | GGCAT 2 |  |  |
| --- | --- | --- | --- | --- | --- | --- | --- | --- | --- | --- | --- |
|  | Time | Mem | Time<br>(8 pcs) | Mem<br>(8 pcs) | Final<br>disk | Time | Mem | Final<br>disk | Time | Mem | Final<br>disk |
| salmonella-128 | 0.22 | 0.47 | 0.40 | 0.41 | 0.00 | 0.20 | **0.24** | 0.01 | **0.11** | 0.69 | **0.00** |
| salmonella-256 | 0.49 | 0.78 | 0.72 | 0.77 | 0.00 | 0.50 | **0.35** | 0.03 | **0.15** | 1.03 | **0.00** |
| salmonella-512 | 1.10 | 1.46 | 1.61 | 1.42 | 0.04 | 1.49 | **0.58** | 0.09 | **0.23** | 1.50 | **0.01** |
| salmonella-1024 | 2.32 | 1.70 | 3.33 | 1.67 | 0.11 | 3.52 | **0.87** | 0.21 | **0.39** | 2.43 | **0.03** |
| salmonella-2048 | 5.30 | 2.64 | 7.58 | 2.38 | 0.33 | 8.83 | **1.56** | 0.51 | **0.76** | 4.10 | **0.09** |
| salmonella-4096 | 10.88 | 2.93 | 15.51 | **2.51** | 0.86 | 22.57 | 3.06 | 1.27 | **1.44** | 6.37 | **0.24** |
| salmonella-8192 | 21.48 | 4.58 | 31.06 | **3.37** | 2.13 | 57.45 | 5.73 | 2.76 | **2.48** | 12.33 | **0.52** |
| salmonella-16384 | 50.10 | 11.26 | 76.82 | **6.38** | 6.46 | 172.74 | 16.32 | 8.28 | **6.09** | 22.41 | **2.18** |
| salmonella-32768 | 131.68 | 24.13 | 202.63 | **10.13** | 16.45 | 435.19 | 34.88 | 15.12 | **13.69** | 29.69 | **10.53** |
| salmonella-65536 | 286.90 | 47.59 | 437.45 | **13.74** | 39.71 | 1067.32 | 71.87 | 34.67 | **33.88** | 47.26 | **31.11** |
| random-128 | 0.50 | 3.57 | 1.07 | 3.52 | 0.00 | 0.74 | 1.64 | 0.10 | **0.14** | **1.41** | **0.00** |
| random-256 | 0.95 | 4.68 | 1.53 | 5.20 | 0.02 | 1.29 | 2.36 | 0.21 | **0.19** | **1.74** | **0.01** |
| random-512 | 2.02 | 7.13 | 3.25 | 7.72 | 0.07 | 2.68 | 4.03 | 0.50 | **0.30** | **2.49** | **0.03** |
| random-1024 | 4.27 | 7.36 | 6.54 | 7.66 | 0.21 | 5.73 | 6.78 | 1.02 | **0.55** | **3.64** | **0.08** |
| random-2048 | 8.48 | 9.06 | 12.57 | 8.86 | 0.62 | 11.96 | 10.82 | 1.73 | **0.95** | **4.98** | **0.21** |
| random-4096 | 17.33 | 11.52 | 25.38 | 10.56 | 1.72 | 26.19 | 18.47 | 3.79 | **1.68** | **7.04** | **0.55** |
| random-8192 | 36.76 | 17.47 | 53.19 | 14.42 | 4.58 | 60.17 | 33.25 | 8.55 | **3.15** | **11.50** | **1.38** |
| random-16384 | 80.90 | 29.57 | 114.33 | 21.46 | 12.41 | 148.18 | 60.28 | 17.65 | **6.04** | **19.07** | **3.61** |

**Figure 4:**
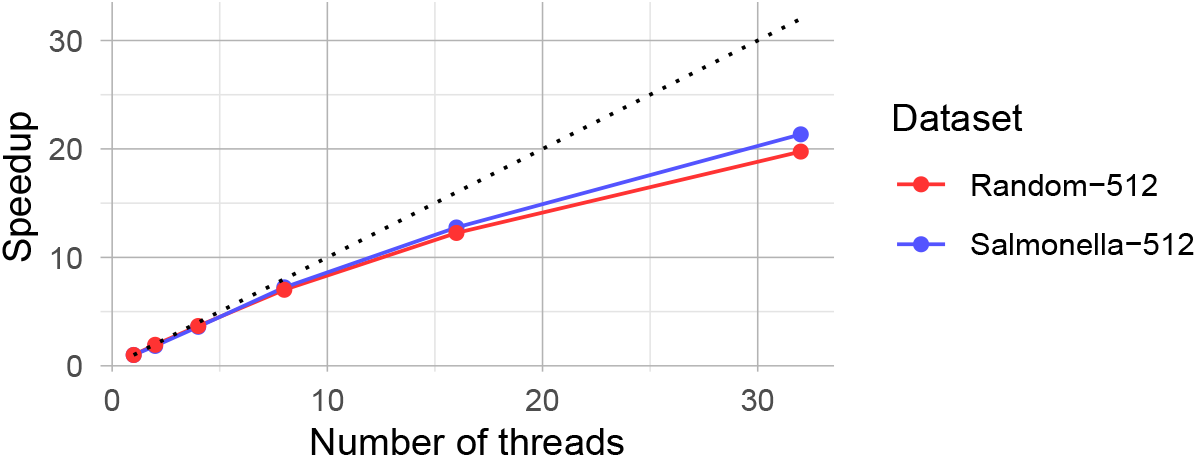
The speedup achieved for index construction on 512 genomes of the random and Salmonella data sets as the number threads are increased. *k* = 31.

#### 4.4.1 Salmonella

In this dataset, the size of the color set structure dominates the size of the *k*-mer indexing structure. We find that GGCAT 2 is faster than both our approach and Bifrost by an order of magnitude. In absolute terms, the time to run our method is still reasonable: On the full dataset, Phases 1, 2 and 3 of our method took 20 min, 132 min and 132 min respectively when run in memory, and Phase 3 still takes only 280 min when run to disk. On the full Salmonella dataset, when the final index is constructed directly to disk, GGCAT uses more RAM than our method on every input size, and 3.4 times more on the full dataset. Despite the three tools using significantly different color set representations, the final disk sizes are similar: 40GB for our sparse-dense index and the SBWT-based *k*-mer index, 35GB for Bifrost and 31GB for GGCAT 2. The construction space overhead, that is, *m*/*d*−1, where *m* is the peak memory and *d* is final disk size, is 20% for our fully in-memory method, 242% for Bifrost and 52% for GGCAT 2. When the final index is constructed directly to disk, our overall peak RAM is only roughly one third of the final index size.

The mean unitig length decreases as the number of unitigs grows, stabilizing at around 48 base pairs (see Table 2). Since the last *k*-mer of every unitig is always marked as a key *k*-mer, we have to mark at least 1/(48 − *k* + 1) ≈5.6% of the *k*-mers as key. We end up marking only a slightly higher amount: a total of 5.8% of all *k*-mers. Overall, the fraction of key *k*-mers seems to be quite well approximated by the fraction of unitig-ending *k*-mers 1/(*L* − *k* + 1), where *L* is the mean unitig length (see Figure 5).

**Table 2:** Dataset statistics for *n* ≥ 128 genomes. The Salmonella datasets are labeled S-*n* and the random datasets are labeled R-*n*. The number of *k*-mers includes both DNA strands, so it is twice the number of canonical *k*-mers. The unitigs statistics are computed from the unidirected de Bruijn graph containing both DNA strands. The mean distinct set size is the average number of elements in the distinct color sets, and the mean color set size is the expected value of the size of the color set of a randomly chosen *k*-mer from the distinct *k*-mers.

| Dataset | Distinct<br>$k$-mers | Key<br>$k$-mers | Distinct<br>color sets | Dense<br>color sets | Sparse<br>color sets | Mean<br>distinct<br>set size | Mean<br>color<br>set size | Number<br>of unitigs | Mean<br>unitig<br>length |
| --- | --- | --- | --- | --- | --- | --- | --- | --- | --- |
| S-128 | 25,820,642 | 2.93 % | 18,415 | 2,593 | 15,822 | 57.96 | 47.68 | 737,677 | 65.00 |
| S-256 | 37,228,100 | 3.75 % | 125,840 | 14,059 | 111,781 | 141.79 | 66.57 | 1,372,922 | 57.12 |
| S-512 | 60,223,428 | 4.61 % | 687,475 | 130,209 | 557,266 | 268.83 | 82.04 | 2,734,306 | 52.03 |
| S-1024 | 79,681,694 | 4.88 % | 1,098,310 | 292,619 | 805,691 | 518.38 | 123.65 | 3,835,925 | 50.77 |
| S-2048 | 123,385,316 | 4.90 % | 1,743,323 | 535,700 | 1,207,623 | 964.60 | 160.11 | 5,925,366 | 50.82 |
| S-4096 | 170,807,566 | 5.21 % | 2,540,284 | 951,462 | 1,588,822 | 1743.32 | 229.74 | 8,686,367 | 49.66 |
| S-8192 | 219,025,616 | 5.31 % | 3,385,675 | 1,410,479 | 1,975,196 | 3217.66 | 357.85 | 11,241,222 | 49.48 |
| S-16384 | 355,596,480 | 5.76 % | 5,841,342 | 2,862,855 | 2,978,487 | 5512.34 | 440.83 | 19,737,346 | 48.02 |
| S-32768 | 514,359,252 | 5.70 % | 8,396,262 | 4,616,836 | 3,779,426 | 9367.01 | 609.37 | 28,289,327 | 48.18 |
| S-65536 | 679,565,754 | 5.80 % | 11,470,379 | 6,992,637 | 4,477,742 | 15911.33 | 921.01 | 37,961,602 | 47.90 |
| R-128 | 290,548,286 | 1.69 % | 188,580 | 50,749 | 137,831 | 15.63 | 3.64 | 4,876,243 | 89.58 |
| R-256 | 425,718,950 | 1.88 % | 706,316 | 330,717 | 375,599 | 26.62 | 4.83 | 7,918,143 | 83.76 |
| R-512 | 749,380,340 | 1.92 % | 1,562,367 | 894,982 | 667,385 | 47.60 | 5.50 | 14,239,936 | 82.63 |
| R-1024 | 1,266,299,900 | 1.98 % | 3,062,665 | 1,975,597 | 1,087,068 | 85.43 | 6.60 | 24,752,934 | 81.16 |
| R-2048 | 1,923,902,914 | 2.07 % | 5,414,584 | 3,874,407 | 1,540,177 | 136.79 | 8.63 | 39,455,816 | 78.76 |
| R-4096 | 3,016,705,654 | 2.24 % | 8,842,659 | 6,816,089 | 2,026,570 | 222.31 | 10.97 | 66,920,361 | 75.08 |
| R-8192 | 4,889,061,902 | 2.38 % | 14,556,798 | 11,929,411 | 2,627,387 | 337.43 | 13.57 | 114,873,088 | 72.56 |
| R-16384 | 7,577,410,586 | 2.59 % | 24,896,466 | 21,422,536 | 3,473,930 | 510.51 | 17.47 | 193,979,805 | 69.06 |

**Figure 5:**
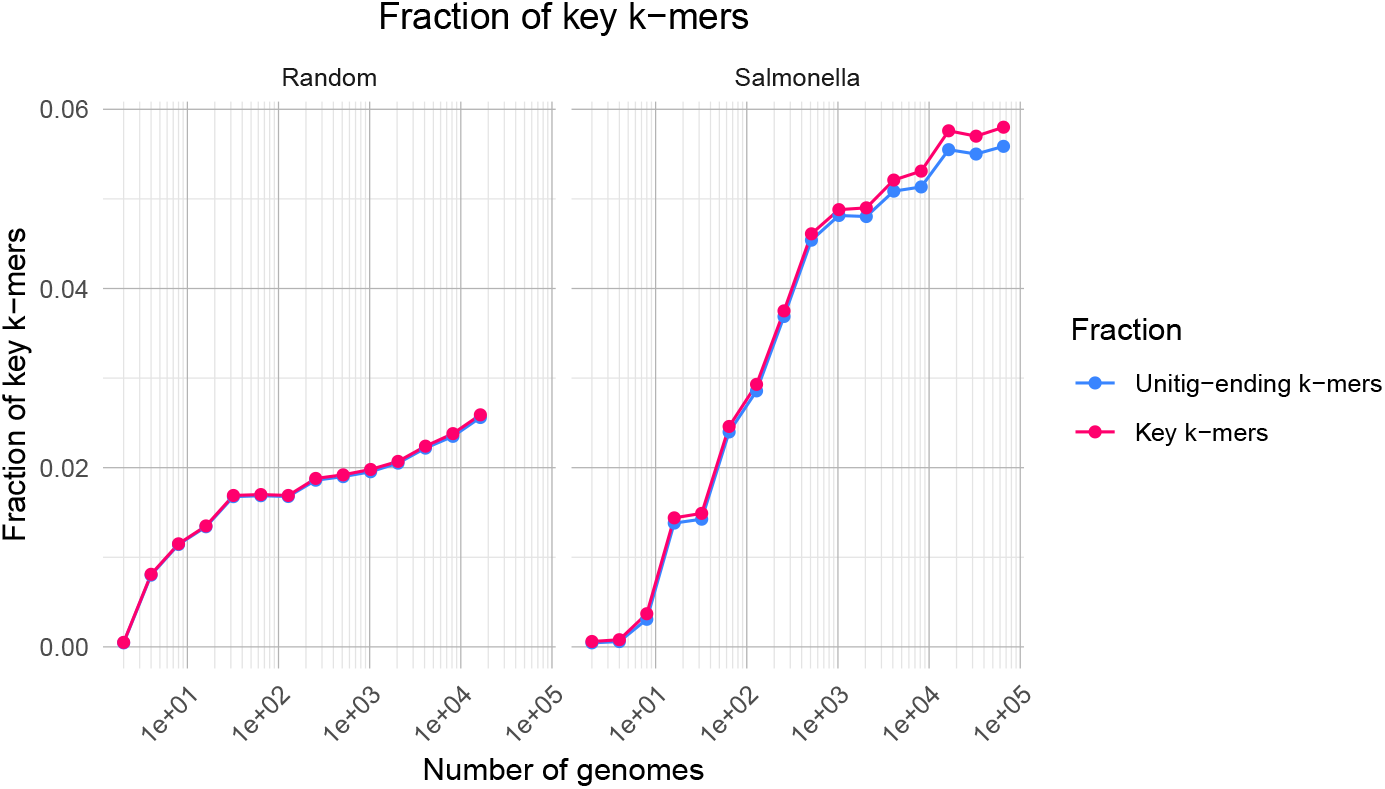
Fraction of *k*-mers (*k* = 31) that are marked as key *k*-mers as a function of the number of genomes. The blue line shows *k*-mers that are at the ends of unitigs, and hence always key *k*-mers.

For a small number of genomes, a vast majority of the distinct color sets are sparse, but at 32,768 genomes, the number of dense sets overtakes the number of sparse sets. In the full dataset, the average distinct color set contains almost one quarter of all colors, making the bitmap representation highly efficient.

#### 4.4.2 Random

In this dataset, the color set structure is small, and the size of the *k*-mer index dominates. The average unitig length was only 69 base pairs on the full dataset. This is consistent with the low number of key *k*-mers, which accounted for only 2.59% of the total *k*-mers, compared to 5.80% in the largest Salmonella dataset. However, this did not translate into reduced space compared to the Salmonella subsets with the same number of genomes, because, even though the sparse-dense representation of the distinct color sets is smaller, the space peak during fingerprinting is much higher.

GGCAT 2 was the best in all the three metrics of time, memory and final disk space. However, in terms of space, our method comes a close second, consuming 21 GiB RAM when the final index is constructed directly to disk, whereas GGCAT takes 19 GiB. For our method, phases 1, 2 and 3 took 11 min, 36 min and 33 min respectively. When writing the final index directly to disk, the time of phase 3 increases to 67 min.

The trends in Figure 6 suggest that if one more data point was added, our method might use less space than GGCAT 2.

**Figure 6:**
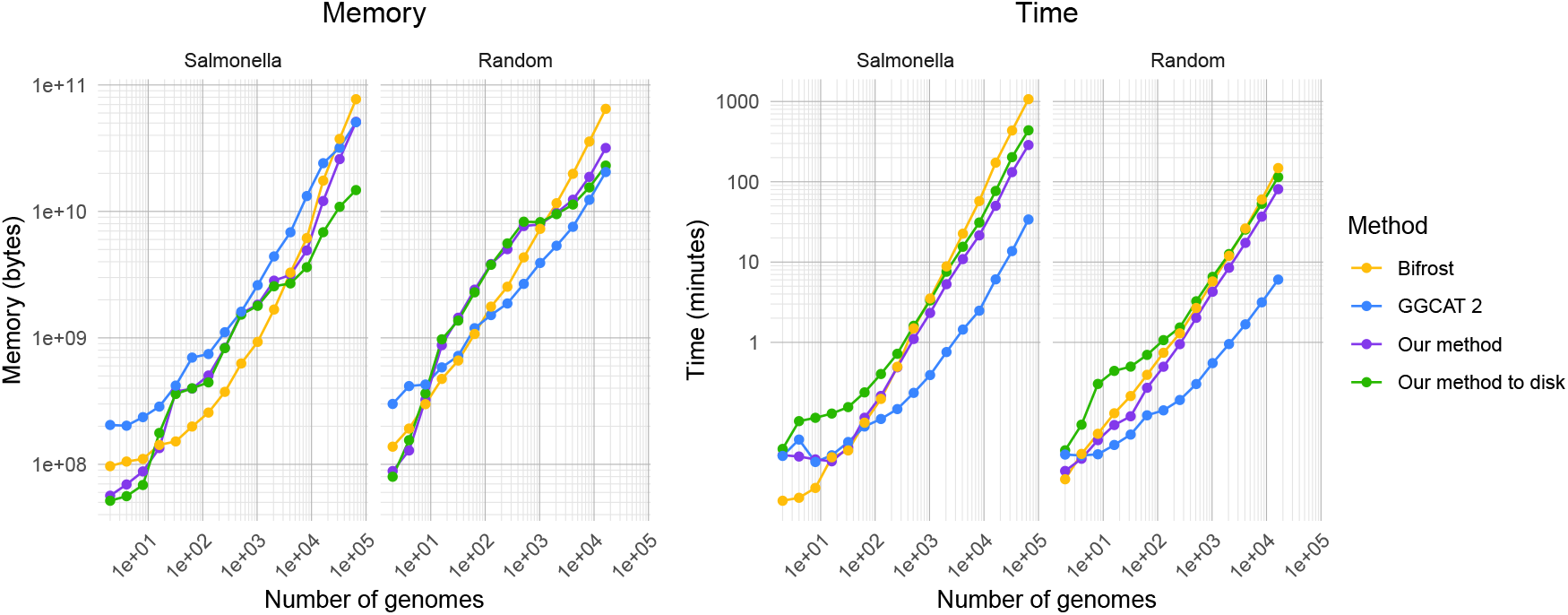
Memory usage (in bytes) and runtime (in minutes) for various competing colored de Bruijn graph construction methods measured in our experiments. *k* is set to 31. Note the double-log scale.

## 5 Concluding Remarks

Given the exploratory nature of genomics, indexing is a frequent operation, making more efficient index construction algorithms, such as the one we have described, of continuing interest.

One direction for future work is to make use of better perfect hashing within our construction algorithm. The current state-of-the-art in minimal perfect hashing, PtrHash [13], claims about 21ns to hash an integer key and 2.41 bits per key. This could lead to significant improvements in our context. Another direction concerns running the construction process from unitigs, which would remove the need for the within-color deduplication structure and so might decrease the peak space and/or improve running time. Preprocessing to unitigs is expensive, however, if the data is already in this form, e.g., unitigs from the Logan project [7], this approach may prove interesting.

Finally, beyond index construction, our approach allows efficient *n*-way merging of colored representations, which is of interest for index updates and may allow more efficient *k*-mer set operations in the context of genomic data exploration [10].

## Footnotes

1 As reported by the command `dd if=/dev/zero of=out.bin bs=1MB count=100000 oflag=direct status=progress`.

